# Unsupervised weights selection for optimal transport based dataset integration

**DOI:** 10.1101/2021.05.12.443561

**Authors:** Aziz Fouché, Andrei Zinovyev

## Abstract

A formulation of the dataset integration problem describes the task of aligning two or more empirical distributions sampled from sources of the same kind, so that records of similar object end up close to one another. We propose a variant of the optimal transport (OT)- and Gromov-Wasserstein (GW)-based dataset integration algorithm introduced in SCOT [Demetci *et al*., 2020]. We formulate a constrained quadratic program to adjust sample weights before OT or GW so that weighted point density is close to be uniform over the point cloud, for a given kernel. We test this method with one synthetic and two real-life datasets from single-cell biology. Weights adjustment allows distributions with similar effective supports but different local densities to be reliably integrated, which is not always the case with the original method. This approach is entirely unsupervised, scales well to thousands of samples and does not depend on dimensionality of the ambient space, which makes it efficient for the analysis of single-cell datasets in biology. We provide an open-source implementation of this method in a Python package, `woti`.

## 1 Introduction

### 1.1 Dataset integration

Recent democratization and flourishing of biological assays at the single-cell level raise important challenges in subsequent analysis pipelines [Lähnemann *et al*., 2020]. One of those, known as data integration and comprehensively described in [Argelaguet *et al*., 2021], is of particular interest and comes in several variants, all related to tying data together across different modalities.

*Horizontal integration* describes the problem of merging two or more datasets expressed in a common feature space, each of those containing samples gathered across distinct sources or experiments. Each of these datasets usually contains biases related to acquisition technique, data collection, experimental variation, preprocessing as well as other factors. In some cases, there can even be dataset-specific cell types; in such case, the integration procedure should ideally detect and avoid to align these dataset-specific instances, called “over-correction”. Horizontal dataset integration techniques have several applications. First, they can be used to construct larger datasets by aggregating a collection of smaller ones, generated across multiple sources or technologies. Simple approaches such as matrix concatenation struggle to get rid of dataset biases, compromising the subsequent use of visualization, clustering, dimensionality reduction and prediction techniques.

*Vertical integration* is the symmetric task of finding a correspondence between matched or unmatched samples from two or more datasets, expressed in different representation spaces. A typical example is with SNARE-seq data [Chen *et al*., 2019], where cells are simultaneously profiled through RNA-seq and ATAC-seq. A common strategy is to perform integration in a latent space in which both representation can be mapped, for instance via matrix factorization techniques [Cantini *et al*., 2019]. Then, vertical dataset integration reduces to horizontal dataset integration in this latent space. The extra layer of difficulty in this approach comes from constructing a relevant latent space via mappings that preserve enough information. Vertical dataset integration is typically used to learn cell types across modalities. If mappings to the latent space are invertible, vertical integration can be used to translate object representations from a domain to another.

A number of approaches have been proposed to solve horizontal and vertical data integration problems. In single-cell biology, mutual nearest neighbors-based methods have become popular [Adey, 2019; Barkas *et al*., 2019]. There also exists techniques based on generative adversarial networks like MAGAN [Amodio and Krishnaswamy, 2018] or variational autoencoders [Simidjievski *et al*., 2019]. We will focus on a class of approaches based on integral probability metrics like MMD-MA [Liu *et al*., 2019] which uses maximum mean discrepancy, or methods using optimal transport (OT). These OT-based methods are generally derived from a color transfer algorithm [Ferradans *et al*., 2013], recently brought into the single-cell field with SCOT [Demetci *et al*., 2020]. We propose a variant of this algorithm, more robust to differences in local density. A comprehensive overview of the available methods is given in [Argelaguet *et al*., 2021].

### 1.2 Optimal transport

OT problem between discrete distributions can be pictured as follows [Peyré *et al*., 2019]. We are given a set of *n* “ware-houses” {*x*_*i*_}_*i*≤*n*_ and a set of *m* “factories” {*y*_*j*_}_*j*≤*m*_. The cost matrix *C* ∈ (ℝ^+^)^*n*×*m*^ contains pairwise transport costs between warehouses and factories: *C*_*ij*_ is the cost required to move one unit of goods from warehouse *i* to factory *j*. Each warehouse contains an amount of goods *w*_*i*_ ∈ ℝ^+^, and each factory requires a quantity of goods *v*_*j*_ ∈ ℝ^+^, we assume 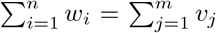. A transport plan between warehouses and factories is uniquely defined by a matrix *P* ∈ (ℝ^+^)^*n*×*m*^, where *P*_*ij*_ is the quantity of goods sent from warehouse *i* to factory *j*. A plan is said to be valid if 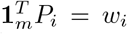 and 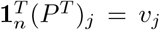 for all *i*≤ *n, j* ≤*m*. The total cost of a transport plan is then defined as

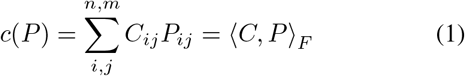

where ⟨.,. ⟩ *F* denotes the Froebenius inner product. A transport plan *P*\* is said to be *optimal* if it is valid and minimizes the cost defined in Eq. 1 over all valid plans.

If factories represent the *reference* distribution and ware-houses the *source* one, we get an intuition behind OT-based dataset integration: a transport plan describes a way to displace the whole mass from the source distribution onto the reference one. The OT plan intuitively favors a natural displacement, with well-preserved local topology as trajectory of masses will typically not cross. Nonetheless, the method is very prone to overfitting, as it can align any distribution onto any other one even if there is no relation between the two. Empirical distributions with similar underlying manifold (for instance, two point clouds where points are shaped as a ring) but different local density also tend to be incorrectly integrated, as shown in section 3.

The optimal transport solution can be approximated using an entropic regularizer, and computed efficiently with the help of Sinkhorn’s algorithm [Cuturi, 2013; Peyré *et al*., 2019].

### 1.3 Gromov-Wasserstein problem

In the general case, defining a cost matrix between two datasets may be challenging: for instance, they can live in different spaces, or one can be a rotation of the other. In these cases, the OT approach is in general unusable. The Gromov-Wasserstein (GW) problem is a natural extension of OT that can overcome these limitations, at the cost of extra hypotheses and computation time. Let 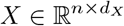 and 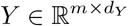 be two datasets living in two metric spaces, and *C*^*X*^ ∈ ℝ^*n*×*n*^ (resp. *C*^*Y*^∈ ℝ^*m*×*m*^) containing pairwise distances between points in *X* (resp. *Y*).

For a transport plan *P*, we define the transport cost with respect to *P* as in [Peyré *et al*., 2019],

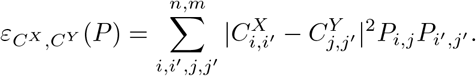

We can get an intuition of why this cost is relevant. The only way to make it small is to find a transport plan *P* so that *P*_*i,j*_ is large if and only if for all *i*′, *j*′, if *P*_*i′*_,_*j′*_ is large then the distance between *x*_*i*_ and *x*_*i′*_ is close to the distance between *y*_*j*_ and *y*_*j′*_. Given two histograms *υ* ∈ ℝ^*n*^ and *w* ∈ ℝ^*m*^, the weighted Gromov-Wasserstein distance between *X* and *Y* is then defined as

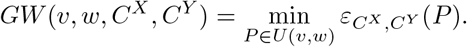

This problem is non-convex in this form, but can be rewritten as a quadratic assignment problem [Loiola *et al*., 2007]. It is NP-hard in the general case, but admits an entropic regularization and can be solved quite efficiently.

### 1.4 Contributions

We propose a dataset integration method based on unbalanced OT/GW dataset integration, that adjusts weights *w*_*i*_ and *v*_*j*_ in source and target empirical distributions before applying OT or GW. This allows for correcting local density disparities in cloud points with similar effective support shape but different local density. We challenge our approach against the balanced technique on three pairs of synthetic and biological datasets. We then discuss limitations of the method, and cases when it is appropriate to use it. We provide an implementation of the method in a Python package, `woti` (weighted optimal transport integration).

## 2. Material and methods

Notebooks for carrying out computational analyses and producing figures are available on `woti`’s GitHub ^1^.

### 2.1 Datasets

Our method was compared to SCOT [Demetci *et al*., 2020] on three pairs of datasets of various dimensions *d*: two synthetic datasets (*d* = 3), two single-cell datasets in cell cycle space (*d* = 2) and one SNARE-seq dataset split in two parts, gene expression (*d* > 10^4^) and chromatin accessibility (*d* = 19). Theoretically, dimensionality does not play a role when integrating datasets using OT and GW but in practice, defining a relevant cost between points in high dimensional spaces is highly challenging - and beyond this paper’s scope.

#### Synthetic datasets

We build two 3D synthetic spiraling datasets unbalanced on purpose. A spiraling shape is interesting for several reasons. First, the underlying shape is continuous which is a good stress test for integration methods. Also, the spiraling pattern can fool integration techniques (some “internal” samples are mapped to the “external” regions for reasons of mass availability). Source spiral contains 500 points split in 4 clusters, and reference spiral 1000 points in one cluster. Spirals are then randomly translated in space, and noise is added.

#### Ewing sarcoma single-cell datasets

RNA-seq datasets were gathered from [Aynaud *et al*., 2020] for Ewing sarcoma patient-derived xenografts (PDX), and from [Miller *et al*., 2020] for Ewing sarcoma cell lines. All raw datasets were preprocessed using standard methods as follows. Cells with less than 200 genes expressed, as well as genes expressed in less than 3 cells were discarded. Then, cells with raw counts below 15,000 or above 50,000 or expressing more than 15% mitochondrial genes were taken out. Cell counts were then normalized to 10,000 counts per cell, before being log-transformed by the function log(1 + *x*). The 10,000 genes with higher variance were kept. All datasets were then smoothed by neighborhood averaging using 10 closest neighbors, using 50 components of PCA for the nearest neighbors computation. The G1/S and G2/M scores for each cell were computed using Ewing sarcoma-specific signatures of cell cycle phases [Aynaud *et al*., 2020]. We chose CHLA9 Ewing sarcoma cell line scRNASeq dataset (*n* = 3752) from [Miller *et al*., 2020], as the reference one, because the differences in cell cycle phases comprised the most important source of transcriptomic heterogeneity in this dataset, which was clear in the simplest 2D PCA projection. PDX352 patient-derived xenograft profiled using scRNASeq (*n* = 1937) was chosen as the source dataset among those published in [Aynaud *et al*., 2020], for its high proportion of non-proliferative cells and with a clear cyclic structure corresponding to proliferating cells.

#### Multi-omics SNARE-seq dataset

Chromatin accessibility and gene expression datasets were generated with SNARE-seq [Chen *et al*., 2019] technology, using a mixture of human cell lines (BJ, H1, K562 and GM12878) [Chen *et al*., 2019]. Every cell was analyzed in both of the assays, and is consequently present in both datasets. Chromatin accessibility records were preprocessed using [Chen *et al*., 2019] guidelines including noise reduction then dimensionality reduction using the cisTopic [González-Blas *et al*., 2019] R package, resulting in a 1047 × 19 matrix. RNA-seq data was normalized to one count per cell for appropriate scaling in the latent space. Counts matrix was log-normalized, then the 1000 top variable genes were kept. Matrix was then Z-score-normalized, and divided by 100. Eventually, a 19-components PCA was carried out in order to match chromatin accessibility dimension. After these steps, datasets were randomly unbalanced by selecting at random a fraction of samples from each cluster. In the chromatin accessibility data, 20% of cells are taken out from cluster 0, 60% from cluster 1 and 80% from cluster 2. In the gene expression dataset, 80% of cells are taken out from cluster 0, 20% from cluster 1 and 60% from cluster 2.

### 2.2 Kernel density uniformization

We propose a density uniformization method to adjust sample weights before unbalanced OT or GW so that, for a given kernel, empirical point density variance is close to be constant over the distribution. Let *X* = {*x*_*i*_}_*i*≤*n*_ be a dataset consisting of *n* samples in a space *𝒳* endowed with a positive semi-definite kernel 𝒦 : *𝒳* × *𝒳* → ℝ^+^. Let us further assume for all *i* ≤ *n*, ∫_*𝒳*_ 𝒦(*x*_*i*_, *x*)*dx* = 1. For every *x* ∈ *𝒳*, we define the empirical point density at *x* as

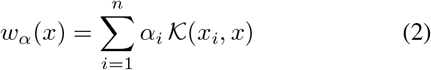

with *α* ∈ ℝ^*n*^. We constraint *α* to the probability simplex so that 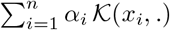. is the PDF of a distribution, meaning *α ⪰* 0 (coordinate-wise comparison) and *α*^*T*^ **1**_*n*_ = 1. We can write an expression for the empirical variance of *w*_*α*_ over the dataset,

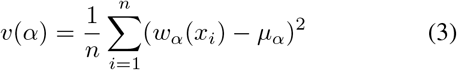

with 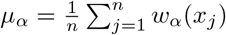.

The kernel uniformization problem can then be stated as minimizing *υ*(*α*) over the probability simplex,

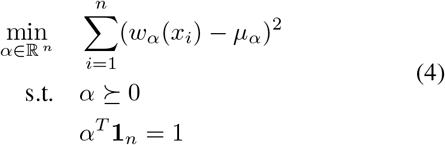

Eq. 3 can be rewritten as a quadratic form,

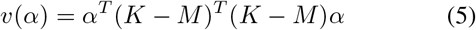

with *K*_*ij*_ = 𝒦(*x*_*j*_, *x*_*i*_) and 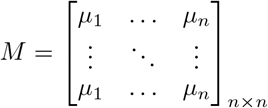 where 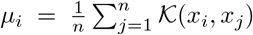.

Eq. 4 defines a quadratic program constrained on a simplex, also known as *standard quadratic optimization problem* [Bomze, 1998] that cannot be solved analytically, motivating the usage of interior point methods. We used the Python implementation of `osqp` [Stellato *et al*., 2020] to carry out the computation, using a Gaussian kernel based on the Euclidean distance matrix (variance-normalized). We reduced the dimension to 3 using PCA for the 19-dimensional SNARE-seq dataset because of the bad scaling of distance-based kernels to higher dimensions. All computation times have been recorded on a desktop computer running Arch Linux, equipped with a 12/24 cores Ryzen 9 3900x, 32GB of DDR4 RAM. Computation was not GPU-accelerated.

### 2.3 Weighted dataset integration

OT or GW can in the discrete case be used as an integration technique for vectorized datasets, originally proposed for histogram color transfer in image processing [Ferradans *et al*., 2013]. Let *X* and *Y* be two matrices of 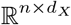 and 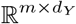 representing two datasets containing respectively *n* and *m* samples. Distance information is necessary to carry out OT or GW. For OT, let *C*_*XY*_ ∈ ℝ^*n*×*m*^ be the matrix containing pairwise distances between *X* and *Y*. For GW, let *C*_*X*_ ∈ ℝ^*n*×*n*^ and *C*_*Y*_ ∈ ℝ^*m*×*m*^ be two matrices containing pairwise distances in *X* and in *Y*. Let *P* ^*^ be the optimal transport plan from *X* to *Y*, computed either using OT or GW, assuming samples from *X* (resp. *Y*) are associated either to uniform weights, or weights obtained via the uniformization procedure described in subsection 2.2. We denote these weights by *α*_*X*_ and *α*_*Y*_. The idea is then to consider each row in diag(*α*_*X*_)^-1^*P*^*^ as a probability distribution. Namely, 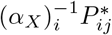 is interpreted as a probability, 𝕡(*x*_*i*_ corresponds to *y*_*j*_). We can then derive an expression for the predicted position of *x*_*i*_ in *Y* representation as the weighted mean point 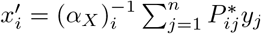, or

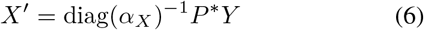

for projecting the whole *X* dataset onto *Y* representation.

OT and GW can be computed with the help of the `Python` package `pot` [Flamary and Courty, 2017]. It notably features C-accelerated implementations of both Wasserstein and GW distances with very good performance, associated to computation times typically between one and ten seconds for all datasets we use.

### 2.4 Assessing integration quality

Assessing integration quality is a non-trivial task, especially when there is no matching between datasets samples. A good integration should not necessarily align all source samples on all reference samples, but rather only align records of similar objects while keeping records of different objects far from one another.

For low dimensional datasets, visual inspection is often sufficient for assessing integration quality, especially when differences are striking between methods. For higher dimension datasets such as SNARE-seq (*d* = 19) though, visual inspection is not reasonable. However, source and reference datasets are *coupled* in the latter case which means each biological cell has been analyzed in both domain, and appears in both datasets. Both dataset contains three well resolved clusters corresponding to different cell lines, that can be separated using a `sklearn` [Pedregosa *et al*., 2011] implementation of the *k*-means algorithm [MacQueen, 1967]. Then, *k*-nearest neighbors model can be trained on the reference dataset, and then predict samples from the integrated source dataset. If integrated clusters are pure and mapped on the right class, the classifier should have a good predictive power. Classifier accuracy is therefore chosen as a first indicator for integration quality.

Though, this evaluation metric present an obvious limitation: it does not take into account how well clusters are merged. In particular, if a cluster *C*_1_ is not merged with its corresponding cluster 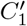 but closer to it than to any other, classifier accuracy will be high but the integration is not achieved. To deal with this issue, we furthermore used the sum of cluster-wise Wasserstein distance (optimal transport cost) as a second integration indicator. It was evaluated in the 3D space where clusters were identified, from integrated *X*′ to reference *Y* datasets, normalized by cluster size *c*_*k*_ with *k* ∈ ⟦1; 3⟧,

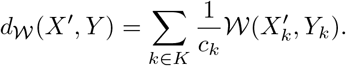

Integration quality can then be represented on a plane where x-axis corresponds to classifier accuracy (higher is better), and y-axis corresponds to cluster-wise Wasserstein distance (lower is better).

## 3. Results

### 3.1 Solving the quadratic program uniformizes density

We demonstrate the effectiveness of density correction on our six datasets. We compare empirical KDE variance over the dataset using equal weights (with *α*_*i*_ = *n*^-1^ for every sample), and using weights minimizing Eq. 4. Tab. 1 shows variance without and with adjusted weights. We see the method decreases KDE variance by orders of magnitude, even if the effectiveness varies from one dataset to another. Computation time varies from 250ms for spiral datasets to 140s for CHLA9. Time is related to dataset size, which conditions the quadratic program dimensionality.

We can visualize an example of density uniformization in Fig. 1. Each cell is represented as a point in a 2D linear subspace of the gene expression space. The x-axis corresponds to the mean expression of G1/S-phase genes and the y-axis corresponding to the expression of G2/M-phase genes. Non-proliferating cells are located on the bottom-left region of the plot, and replicating cells travel around the loop in the anti-clockwise direction.

**Figure 1:**
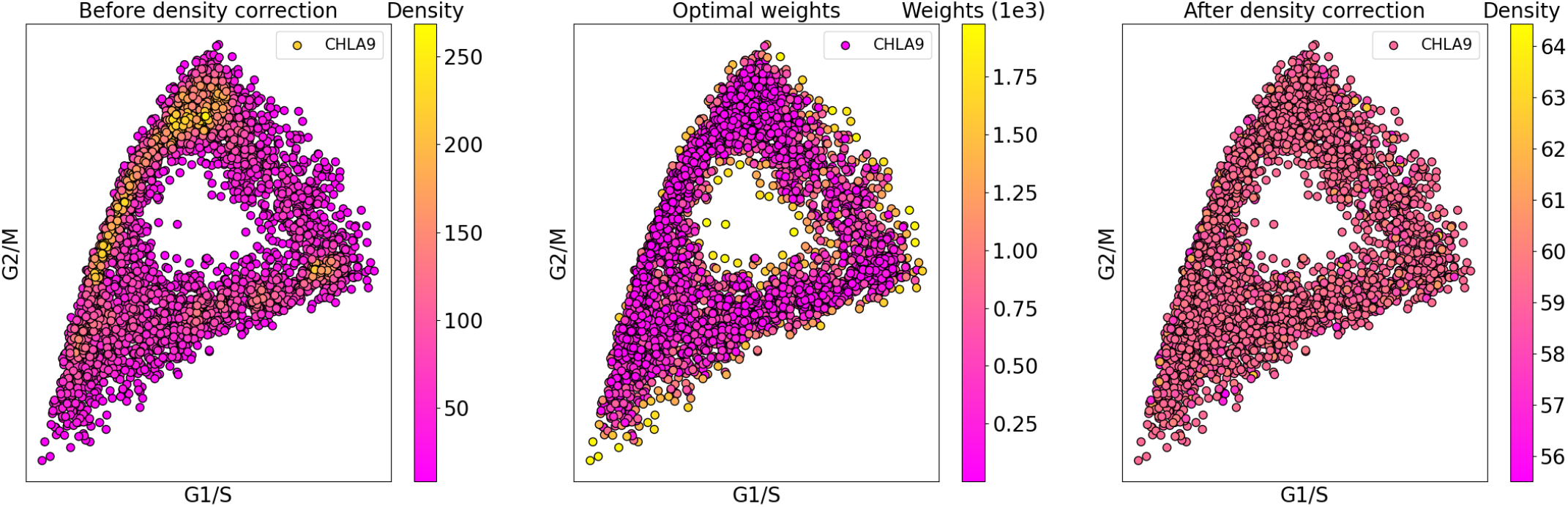
Density correction applied on a single-cell dataset (CHLA9) in G1/S G2/M space. **Left:** KDE with equally weighted kernels. **Middle:** Quadratic program solution. **Right:** KDE with kernels weighted according to the quadratic program solution.

Middle pane of Fig. 1 presents the choice for optimal kernel coefficients in each dataset. The average coefficient is the inverse of dataset size, 2.6 × 10^−4^. As expected, samples in populated regions are associated to below-average coefficients while samples in sparse regions like loop borders are associated to above-average coefficients. This suggests the importance for the reference dataset to be of high quality, without outliers as they would be associated to high coefficients and probably fool downstream methods. Right pane of Fig. 1 shows density-corrected kernel density estimation on each dataset. We can see KDE variability between the nodes is greatly diminished as expected.

**Table 1:** KDE variance over each dataset with equal weights (before) and optimal weights (after). SpA/SpB refers to spiral synthetic datasets (spirals) A and B.

| Dataset | SpA | SpB | PDX | CHLA | RNA | ATAC |
| --- | --- | --- | --- | --- | --- | --- |
| Before | 686 | 2267 | 3956 | 7.79 | 28472 | 258 |
| After | 16 | 37 | 0.16 | 0.003 | 82 | 10 |

### 3.2 Integration improvement on synthetic unbalanced datasets

We first use two synthetic 3D spiraling datasets to assess the effectiveness of weighted OT integration (WOTi) versus balanced OT integration (OTi). Results are presented in Fig. 2. Left-hand plot presents a PCA view of both datasets in their initial state. The reference spiral B (grey) simulates a high quality dataset (uniform distribution along the manifold, high number of samples) while the source spiral A (colored) simulates a low quality dataset (non-uniform repartition along the manifold, fewer cells). A good integration should map the piece-wise spiral onto the reference one in the corresponding regions, preserving color gradient and clusters.

**Figure 2:**
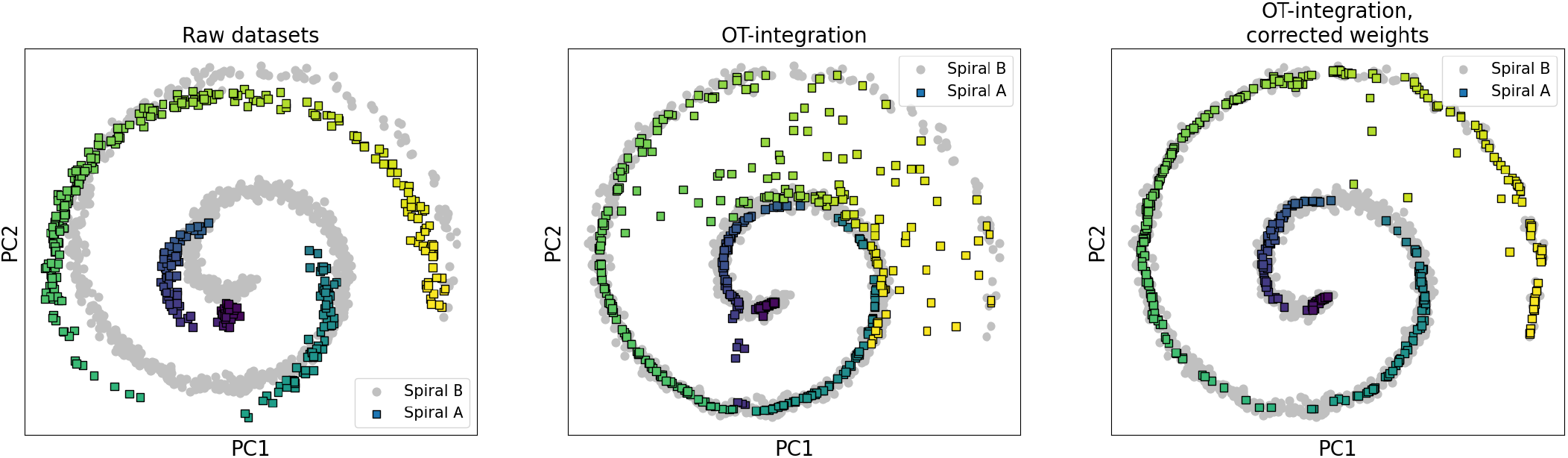
OT based integration of two synthetic 3D datasets (PCA representation), source points are colored based on their initial position in the spiral. **Left:** Raw datasets. **Middle:** OT integration. **Right:** Weighted OT integration.

OTi fails in all these tasks (middle pane). Many points fall outside the reference, yellow and blue points are merged in the center and clusters are indistinguishable. WOTi takes an extra 2s of time for preprocessing weights, but yields a significant improvement in integration quality (right pane). All source points but a dozen fall on the reference distribution, color gradient is respected and the four initial clusters clearly appear. WOTi outperforms OTi on this synthetic example in all aspects, except for the two extra seconds of preprocessing time.

### 3.3 Integration improvement of cell-cycle trajectory in a single-cell dataset

Integrating cells in the cell cycle space has several applications. It can be used to correct the loop shape of an average quality dataset with respect to a high quality reference dataset, or to infer cell cycle state of cells in a semi-supervised fashion with label transfer methods if the reference dataset is labeled. Here, we integrate a Ewing sarcoma PDX352 dataset of 1937 cells with a majority of non-proliferating cells onto a Ewing sarcoma cell lines dataset CHLA9 mainly composed of proliferating cells (Fig 3) A good integration should result on mapping non-proliferating cells (located on the bottom-left of a cell cycle plot, dark blue points) of PDX onto non-proliferating cells of CHLA9, while preserving the distribution of proliferating cells in PDX over the cell cycle loop.

**Figure 3:**
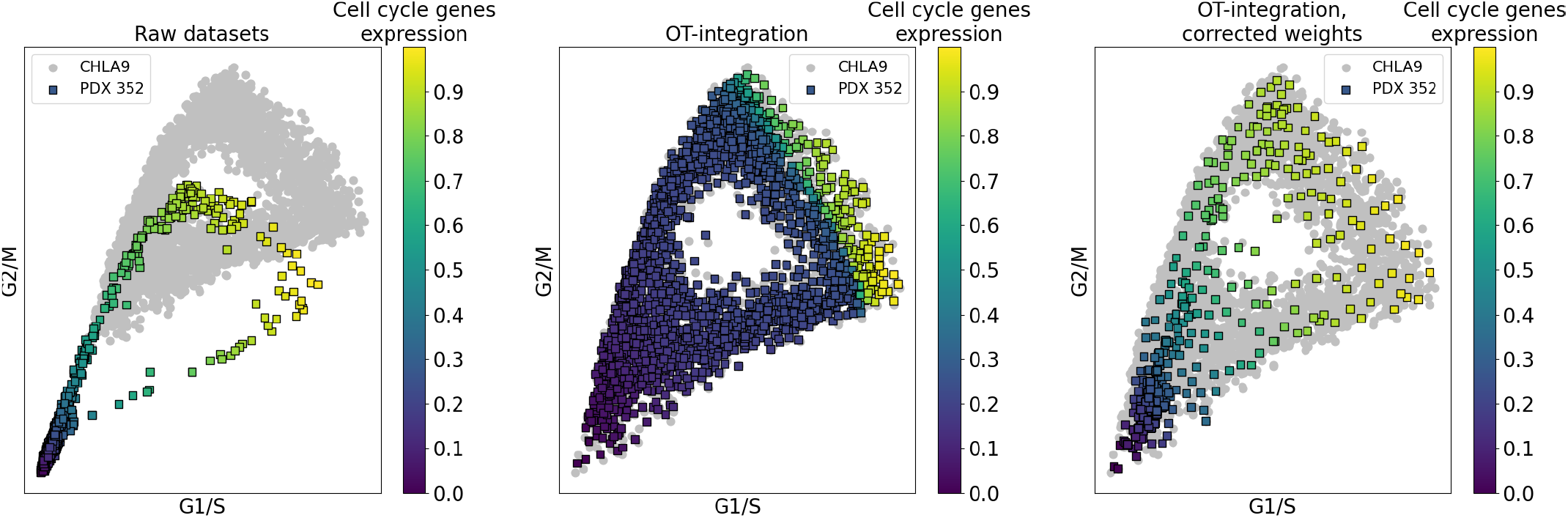
OT based integration of single-cell Ewing sarcoma datasets in the cell cycle space. Cells are colored by the average cell cycle genes expression in PDX dataset, scaled to unity interval. **Left:** Raw datasets. **Middle:** OT integration. **Right:** Weighted OT integration.

Fig. 3 presents integration results. Left-hand pane shows data before integration, the colored dataset being the source PDX352, and the grey dataset being the reference CHLA9 dataset. Middle pane displays integration performed with the state-of-the-art OTi, and integration by WOTi stands on the right-hand pane. Source cells are colored with respect to their position in the initial PDX352 dataset. Dark blue points correspond to non-proliferating cells (low cell cycle genes expression), yellow points to cells expressing cell cycle genes at high level. A good integration technique must maintain the initial loop shape and color pattern, i.e. map proliferating and non-proliferating cells in both datasets. When performing OTi (middle pane), we see the integrated PDX cell cycle loop shape perfectly overlaps with CHLA9 one. We observe most non-proliferative cells (dark blue) are incorrectly projected onto a proliferative state in the cell cycle loop, while they should stay clustered in the bottom-left corner, as a result of high unbalance in distributions between datasets. WOTi (right-hand pane) allows non-proliferating cells to be correctly maintained to the bottom-left part of the plot, and to recover a loop coloring resembling the initial dataset, integrated with cells in CHLA9 dataset. Furthermore, PDX initial loop shape overlaps correctly on CHLA9, with points nicely distributed in all CHLA9 areas and a high density cluster representing non-proliferative cells.

### 3.4 Integration improvement in multi-omics data

Unsupervised vertical dataset integration is a challenging task requiring to map similar samples between two representations. It does not only require a robust alignment technique, but also clever cross-representation projections to construct a meaningful common space for integration to take place. In our multi-omics dataset, performing a 19-components PCA on gene expression data, and reducing chromatin accessibility data to a 19 dimensional space was enough to recover close corresponding clusters - this is not the general case. Both OT integration methods deliver equally good results in the balanced case, with a near-perfect integration (not shown in figures, see Supplementary materials and also results from [Demetci *et al*., 2020]). We challenge the methods by randomly unbalancing clusters in both datasets.

We assess integration quality on Fig. 4, both visually (three first plots) with visual inspection of clusters merging, and quantitatively (fourth plot). In the latter representation, the x-axis represents a *k*-NN classifier accuracy over the source dataset in the task of cluster classification, when trained on the reference dataset (higher is better). The y-axis represents the sum of Wasserstein distances between source and reference clusters (lower is better). The random unbalance procedure has been carried out 100 times, giving these point clouds. OTi strikingly suffers from the unbalance in cluster sizes between data modalities. Orange and purple dots pollute the cyan cluster, and data is considered less integrated with respect to both metrics (Fig. 4, left pane) than the datasets before integration. In the other hand, WOTi yields much better results. Visual inspection reveals cluster structure is well preserved, and points typically overlap well onto the right cluster despite class disequilibrium. Classifier accuracy is similar to the one before integration, while distance between clusters is greatly reduced.

**Figure 4:**
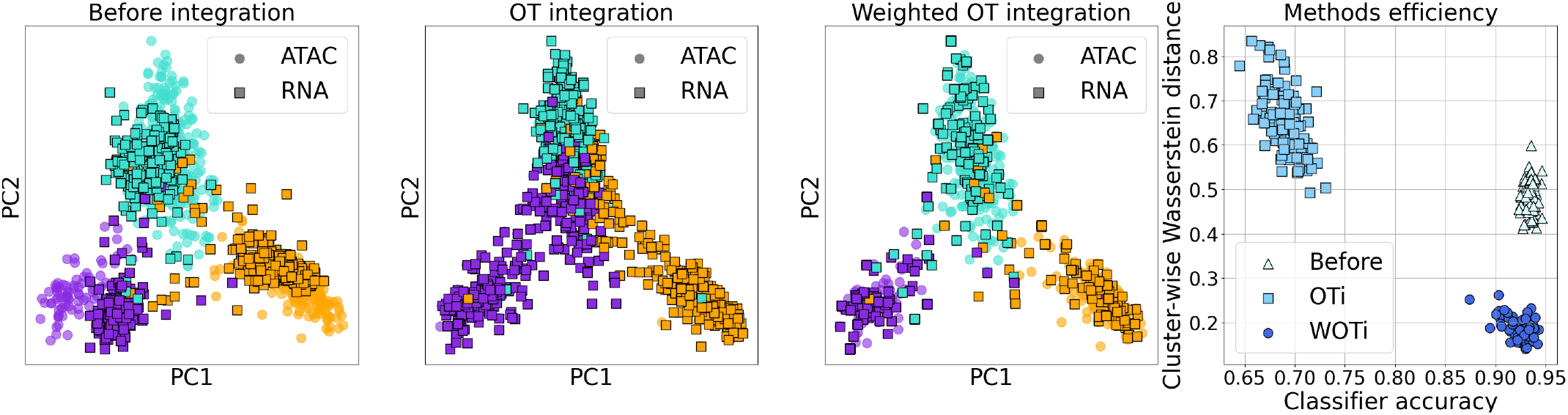
Comparison of both OT integration methods on coupled SNARE-seq data consisting of chromatin accessibility (ATAC-seq) and gene expression (RNA-seq) records, in PC space. Points are colored with depending on which cluster a cell belongs to in ATAC representation. **Left:** Datasets before integration. **Middle left:** OT integration. **Middle right:** Weighted OT integration. **Right:** Performance of optimal transport integration methods with (WOTi) and without (OTi) density correction, on 100 i.i.d. samplings for datasets unbalance.

In this case, building the latent space was quite easy using PCA projections, but can be way harder for other datasets. Once this was done, our method effectively integrates 19D data, generally mapping cells on their corresponding cluster. Computation time for OT is less than 1s for each dataset. Pre-processing the weights takes 2.5s for chromatin data, and 0.9s for RNA-seq data.

## 4 Discussion

We present a variation of the OT dataset integration technique, originally designed for histogram color transfer but finding applications in a variety of domains, notably in single-cell with SCOT. We address the problem of integrating datasets with similar effective support but different local density. These types of datasets are frequent in real-life applications, such as single-cell biology. We propose a way to choose sample weights for the unbalanced OT problem so that dense regions are associated to lower weights and vice-versa. We demonstrate the effectiveness of this approach on three pairs of datasets of various dimensions. In particular, we demonstrate our method outperforms original OT integration for all datasets when local density is unbalanced, with reasonable computation time.

Choosing between OT and GW is a concern that must be addressed in every real-case application. OT is appliable only when a relevant metric can be defined between samples of different datasets, and will typically avoid any kind of transformation, which can help in the case of symmetrical datasets for instance (SNARE-seq is such a case). On the other hand, GW is typically invariant with respect to isometries and can be applied event when both datasets to integrate do not share the same data space.

Though, WOTi does not solve all limitations of OT dataset integration. For instance, it cannot deal with datasets in which only a subset of samples represent objects of the same type. In this case, the method will overfit and align all source records onto the reference ones, even if some of them should not be aligned at all. Using distance-based cost matrices also does not scale well with high dimensional data, so designing a ground cost suitable to high dimension data for performing OT integration is still an open question, recently addressed in [Huizing *et al*., 2021]. Solving a quadratic problem in the probability simplex in a very efficient way is challenging in high dimension, but we did not find this to be a decisive obstacle as applications for which we use WOTi do not exceed 5000 sample points. With such dataset size, standard interior point methods are efficient enough to yield reasonable computation time (from a second to a minute). Alternative formulations and strategies must exist though, to at least approximate the result in a more efficient manner in our particular case. Finally, vertical dataset integration using WOTi highly depends on the latent space construction which is still an unsolved question in general, probably highly dependent on the application field.

The question of outliers is a serious concern when performing density uniformization, not addressed yet. Indeed, it is easy to show that using a variance-normalized distance matrix to define kernel density is not robust to distant outliers. In the limit case, the normalized distance matrix tends to represent a two-points setup with the outlier as one point, and all the other points degenerated into one location. The solution is there to put a weight of 1*/*2 on the outlier, and a weight of 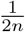 on all the other points. This is of course very detrimental, and must be for now addressed upstream by outlier filtering, using one of the available and computationally tractable methods.

Although, dataset integration using OT is a promising method which can yield high quality results, as with single-cell datasets in the cell cycle space or SNARE-seq multiomics data; we showed choosing weights is a key part of the problem. If making all weights equal is a tempting (and easy) approach, we demonstrate it fails with unbalanced datasets that are usual when dealing with real-life data. Performing balanced OT integration can in these cases lead to severe overfitting. We believe developing algorithms to choose sample weights like in WOTi is a critical step to ensure meaningful dataset integration, and we are looking forward to seeing more approaches to do so.

## 5 Acknowledgements

This work has been partially supported by the French government under management of Agence Nationale de la Recherche as part of the “Investissements d’Avenir” program, reference ANR-19-P3IA-0001 (PRAIRIE 3IA Institute) and by the Ministry of Science and Higher Education of Russian Federation (Project No. 075-15-2020-808). A. F. is supported by Ecole Normale Supérieure Paris-Saclay.

## Footnotes

1 https://github.com/Risitop/WOTi

## Notes

### Competing Interest Statement

The authors have declared no competing interest.

